# IL-2 Modulates Th2 cell Responses to Glucocorticoid: A Cause of Persistent Type 2 Inflammation?

**DOI:** 10.1101/365783

**Authors:** Tharsan Kanagalingam, Meerah Vijeyakumaran, Nami Shrestha Palikhe, Lauren Solomon, Harissios Vliagoftis, Lisa Cameron

## Abstract

**Background:** Inhaled glucocorticosteroids (GCs) are the main treatment for asthma as they reduce type 2 cytokine (IL-4, IL-5 and IL-13) expression and induce apoptosis. Asthma severity is associated with GC insensitivity, increased type 2 inflammation and circulating Th2 cells. Since IL-2 is a T cell survival factor, we assessed whether IL-2 levels associate with the proportion of Th2 cells and/or correlate with clinical features of asthma severity.

**Methods:** Peripheral blood from asthma patients (n=18) was obtained and Th2 cell numbers determined by flow cytometry. Peripheral blood cells were activated with mitogen (24hrs) and supernatant levels of IL-2 and IL-13 measured by ELISA. *In vitro* differentiated Th2 cells were treated with dexamethasone and IL-2 and assessed for apoptosis by flow cytometry staining of Annexin V. Level of mRNA for anti-apoptotic (BCL-2) and pro-apoptotic (BIM) genes as well as IL-13 were determined by qRT-PCR.

**Results:** IL-2 produced by activated peripheral blood cells correlated negatively with lung function (FEV_1_) and positively with daily dose of inhaled GC. When patients were stratified based on IL-2 level, high IL-2 producers made more IL-13 and had more circulating Th2 cells. *In vitro*, increasing the level of IL-2 in the culture media was associated with resistance to DEX-induced apoptosis, more BCL-2 and less BIM mRNA. Th2 cells cultured with higher IL-2 also had more IL-13 mRNA and required higher concentrations of DEX for cytokine suppression.

**Conclusions and Clinical Relevance:** IL-2 modulates Th2 cell responses to GC, supporting both their survival and pro-inflammatory capacity, suggesting that a patient’s potential to produce IL-2 may be a determinant in asthma severity.

## INTRODUCTION

Asthma is a syndrome characterized by symptoms of diverse pathogenesis (1, 2). Type 2 cytokines promote many of the processes responsible for the development of asthma and symptom manifestation. IL-4 is essential for Th2 cell differentiation and drives B cell isotype switching to IgE; IL-5 is a differentiation factor for eosinophils and mediates their egress from the bone marrow during allergic responses; IL-13 mediates airway hyperresponsiveness through inflammatory cell infiltration, smooth muscle contraction and epithelial secretions [reviewed in (3)]. These cytokines are produced by both type 2 CD4+ helper T cells (Th2 cells) (4, 5) and group 2 innate lymphoid cells (ILC2s) (6). ILC2s are innate immune cells found mainly within tissues (7, 8), activated by the epithelial-derived cytokines IL-25 and IL-33 (9). Th2 cells are allergen-specific memory T cells that circulate between the lymph nodes and circulation, infiltrating tissues upon allergen exposure (10-12). While both cell types contribute to overall type 2 cytokine levels, Th2 cells are considered the principle cell population responsible for their rapid release upon allergen re-exposure as well as maintenance of chronic allergic inflammation (11, 13).

Glucocorticoids (GCs) are the main treatment for asthma (14). Their efficacy is considered, in large part, to be due to their ability to suppress type 2 cytokines, *in vivo* (15), *ex vivo* (16) and *in vitro* (17). In most cases inhaled GC therapy is sufficient to achieve asthma control, though the amount required varies greatly amongst patients (18). In severe asthma, however, even high doses of inhaled GC fail to control symptoms and adequately improve poor lung function (18). Studies show that some moderate/severe asthmatics experience improved exacerbation rates, lung function and eosinophilia after using anti-Th2 therapies that block IL-5 and IL-13 (19-22), revealing that persistent type 2 inflammation was a factor in their disease severity.

Another anti-inflammatory effect of GC is their ability to induce apoptosis (23). This has been well demonstrated for primary human eosinophils (24, 25), though for T cells the effects vary based on subset examined. For instance, GCs effectively induce apoptosis of thymocytes (26), while memory T cells are less sensitive (27, 28). Studies with murine T cells indicate Th2 cells are less sensitive than Th1 cells to GC-induced apoptosis (29). Recently, we reported higher levels of serum IL-13 as well as circulating Th2 cells in severe compared to mild/moderate asthmatics (30). Others have shown Th2 are also higher in bronchoalveolar lavage of severe compared to non-severe asthmatics (31). As such, asthma severity is related not only to persistent type 2 cytokine expression but also to the inability of GC therapy to eliminate Th2 cells.

One mechanism that could regulate the sensitivity of Th2 cells to GC is the IL-2 pathway. IL-2 is a T cell growth and survival factor that promotes differentiation of the memory T cell phenotype (32). It signals through the IL-2 receptor (IL-2R) which couples with Janus kinases (JAK)s to activate STAT5 transcription factors (33). Studies in T cell lines have shown that IL-2 interferes with GC receptor (GR) nuclear translocation, reducing signaling (34) and inhibiting GC-induced apoptosis (35). Single nucleotide polymorphisms in IL-2 and the IL-2R have been associated with asthma severity (36), suggesting the strength of the IL-2 pathway could be a factor in T cell resistance to GC. To investigate this, we examined IL-2 production from peripheral blood cells and its relationship with clinical features of asthma, type 2 inflammation and the effect of this cytokine on GC responses of human Th2 cells.

## METHODS

### Subjects

This study was approved by the institutional review board of the University of Alberta (approval number PRO1784). All subjects gave informed consent. Patients were recruited from the tertiary care Asthma Clinic at the University of Alberta, where all clinical measures were obtained. Severe asthma was defined as patients on high dose inhaled corticosteroids (ICS, ≥ 1000 μg/day fluticasone equivalent), a 2^nd^ line controller (long-acting beta agonist, leukotriene modifier/or theophylline) and/or oral corticosteroid (OCS) therapy for ≥ 50% of the previous year who remain uncontrolled despite this therapy (18).

### Peripheral blood cell activation and IL-2 measurement

Whole blood (1ml) from asthmatic patients was obtained, diluted 1:1 with RPMI 1640 and activated with mitogen (PMA, 20ng/ml; ionomycin, 1μM) for 24 hours at 37°C. IL-2 and IL-13 levels were measured in the supernatants with IL-2 (R&D Systems, DY202-05) and IL-13 (Diaclone, 851.630.005) ELISA. Lower limit of detection for IL-2 was 1.7pg/mL and IL-13 was 3.1 pg/mL.

### Flow Cytometry

#### Profiling of peripheral blood cells

The proportion of helper T cells and Th2 cells in peripheral blood were identified by whole blood staining as in (30). In brief, antibodies against CD4 (Clone 1F6; Serotec, Oxford, UK) and CRTh2 (clone BM16; Miltenyi Biotech) were used to determine the proportion of cells exhibiting positive staining for these markers, as assessed by flow cytometry. The proportion of helper T cells was identified by low side scatter (SSC^low^) and high CD4^(high)^. Th2 cells were (SSC^low^), CD4^(high)^ and CRTh2 positive and reported as a proportion of total white blood cells (WBC). Flow cytometry data were collected on BD LSR Fortessa (BD, CA, USA) using FACS Diva software. Gates were set in accordance with the profiles of the isotype control and/or negative control beads.

#### Apoptosis

Both primary Th2 cells and CCRF-CEM cells were assessed for apoptosis and cell death using Annexin V and 7-AAD staining (Biolegend, CA, USA). Cells (0.5 × 10^6^) were washed with 1mL of PBS flow cytometry activated cell sorting (FACS) buffer (0.5% bovine serum albumin, 0.1% sodium azide, 3% FBS) then pelleted at 4°C. Cells were resuspend in 100μL of Annexin V binding buffer (10mM HEPES pH 7.4, 140mM NaCl, 2.5mM CaCl_2_), and stained with 1μL of Alexa Fluor 647 Annexin V (BioLegend-640911, CA, USA) and 5μL of 7-AAD Viability Staining Solution (BioLegend, CA, USA). Data were acquired using an LSR II (BD Biosciences) and analyzed with Flowjo (Version 10, Ashland, OR, USA). Data are reported as the proportion of the total cell population.

### Cell culture

#### Primary Th2 cells

Peripheral blood mononuclear cells (PBMC) from healthy donors were obtained by density centrifugation using Ficoll Histopaque Plus (GE Healthcare, Sweden). As previously described (37), helper T cells were isolated by negative selection with a CD4+ T cell Isolation Kit II (Miltenyi Biotech, CA, USA) and cultured in X-Vivo 15 media (Lonza, MD, USA) supplemented with 10% Fetal Bovine Serum (Wisent, QC, Canada), 1X Penicillin-Streptomycin-Glutamine (Gibco, ON, Canada). For Th2 differentiation, CD4+ T cells were activated for 3 days on plate bound antibody (α) against CD3 (clone UCHT1, 1μg/mL) and αCD28 (clone 37407, 1μg/mL) in the presence of recombinant human rhIL-2 (5ng/mL, R&D Systems, MN, USA), rhIL-4 (10-20ng/mL, R&D Systems, MN, USA), blocking antibody for IFNγ (AF-285-NA, 1μg/mL) and IL-12 (Clone C8.6, 1μg/mL). After activation, cells were proliferated with the same concentration of cytokines (IL-2 and IL-4) and blocking antibodies (IFNγ and IL-12) for 4 days. Following 2 rounds of this differentiation protocol, Th2 cells (CRTh2^+^ CD4^+^, purity of ~98%) were obtained using a CRTh2 isolation kit (Miltenyi Biotech, CA, USA). Th2 cells were maintained on cycles of IL-2 and plate bound αCD3/αCD28 (3 days) or just IL-2 (4 days) at 2×10^6^ cells/mL. For experiments, Th2 cells (1.3 × 10^6^/ml) were examined following exposure to various concentrations of dexamethasone (DEX, Sigma Aldrich, ON, Canada) and IL-2.

#### Immortalized Tcells

CCRF-CEM cells (clone CRM-CCL-119) are a CD4+ T cell line derived from lymphocytic leukemia and were purchased from American Type Culture Collection (VA, USA). Cells were grown in RPMI-1640 media (Sigma Aldrich, ON, Canada) supplemented with 10% Fetal Bovine Serum (Hyclone Scientific, Fisher Scientific, Ontario, Canada) and 1X Penicillin-Streptomycin-Glutamine (Gibco, ON, Canada). Cells were incubated at 37°C, in 85% humidity and 5% CO2. Cells were maintained at 0.2 × 10^6^ cells/mL and were re-seeded every 2 days.

### Quantitative real time polymerase chain reaction (qRT-PCR)

Messenger RNA was isolated using the RNeasy mini plus extraction kit (Qiagen, ON, Canada) and cDNA was synthesized from 400 ng RNA using iScript reverse transcriptase (BioRad, CA, USA). Taqman gene expression assays (Life Technologies, CA, USA) for IL-13 (Hs00174379_m 1), BCL-2 (Hs00608023_m1), BIM/BCL-2L11 (Hs00708019_s1) and GAPDH (Hs02786624_g1) were used. Data were analyzed using cycle threshold (Ct) relative to housekeeping gene GAPDH. Fold increase relative to control condition was assessed for experimental treatments using 2^−ΔΔCt^.

### Statistical analysis

Patient data stratified by IL-2 levels were analyzed for statistical significance using Pearson’s Chi-Square test for categorical data and Independent *t*-test for continuous variables. For cell culture experiments, statistical significance for apoptosis and gene expression were determined by ANOVA with post hoc analysis (Student-Newman-Keuls method) or Student’s t-test. Correlations were determined using Pearson’s Correlation. Data was analyzed using SigmaPlot Version 12.5 and considered significant with *p* < 0.05.

## RESULTS

### Peripheral blood cell production of IL-2 associates with asthma severity

Patients were recruited (n=18) from new referrals to our Asthma Center and clinically characterized to assess asthma severity. The amount of IL-2 produced by activated peripheral blood cells from these patients was highly variable, ranging from 14 − 102,933 pg/mL. Patients stratified based on median IL-2 production (42,600 pg/ml) showed no difference in age, body mass index, IgE levels or smoking status. However, those with high IL-2 (> median) did have lower FEV1 and were taking a higher daily dose of inhaled corticosteroid (Table 1). Analyzing the whole population together, IL-2 production was inversely correlated with FEV1 (r = − 0.558, *p* = 0.0162; Fig. 1A) and positively correlated with total daily dose of inhaled corticosteroid (r = 0.561, *p* = 0.0155; Fig. 1B), suggesting its association with clinical features of asthma severity. While this study population consisted of only 4 severe asthmatics, our analysis does suggest this group is characterized by higher IL-2 production (56,380 pg/ml) compared to non-severe asthmatics (32,133 pg/ml, *p* = 0.07).

**Table 1.**

| IL-2 (pg/ml) | < 42,600* > |  | P -value |
| --- | --- | --- | --- |
| Number | 9 | 9 |  |
| IL-2 (mean, pg/ml) | 14,221 | 60,822 |  |
| Sex (% F) | 67% | 44% | ns |
| Age | 42.1 | 38.9 | ns |
| Body Mass Index (BMI) | 31.97 | 37.76 | ns |
| IgE (kU/L) | 1.86 | 0.16 | ns |
| History of Smoking (%) | 44.4 (4/9) | 44.4 (4/9) | ns |
| Pack year | 2.94 | 7.88 | ns |
| Currently Smoking (%) | 0 | 0 | ns |
| FEV<sub>1</sub><sup>#</sup> (% Predicted) | 87.6 | 74.3 | **0.047** |
| FEV<sub>1</sub>/FVC<sup>†</sup> | 71.22 | 64.77 | ns |
| Total daily dose ICS<sup>‡</sup> (µg/day) | 293.8 | 722.3 | **0.026** |
| Severe Asthma (%)<sup>§</sup> | 11 (1/9) | 22 (3/9) | ns |
| Eosinophils (% complete blood count) | 3.267 | 3 | 0.823 |
| % CD4<sup>+</sup> T cells | 4.54 | 7.98 | 0.070 |
| % CD4<sup>+</sup> CRTh2<sup>+</sup> | 4.687 | 4.652 | ns |
| % CD4<sup>+</sup> CRTh2<sup>+</sup>/WBC<sup>&</sup> | 0.167 | 0.353 | **0.005** |
| IL-13 (pg/ml) | 306.8 | 584 | **0.042** |

\* Data stratified by median IL-2 (42,600 pg/ml); <sup>#</sup> Forced expiratory volume in one second
<sup>†</sup> Forced vital capacity; <sup>‡</sup> ICS, inhaled corticosteroid, fluticasone equivalent
<sup>§</sup> ATS/ERS Guidelines<sup>20</sup>; <sup>&</sup> WBC, peripheral white blood cells

**Figure 1.**
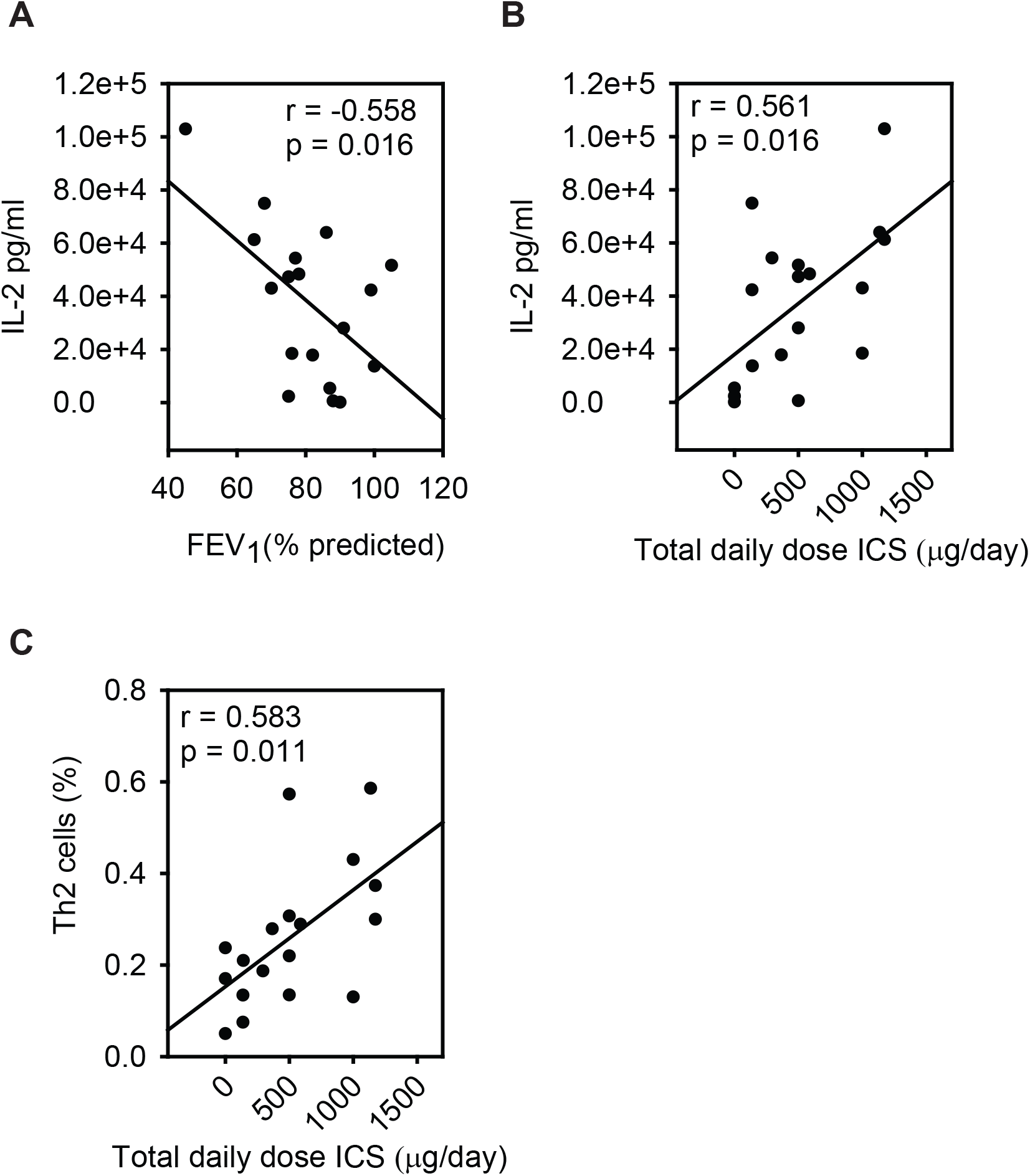
*IL-2 production associates with asthma severity **A)*** IL-2 production was inversely correlated with FEV1 (% predicted) and ***B)*** positively correlated with total daily dose of inhaled corticosteroid (fluticasone equivalent μg/day). ***C)*** Percentage of circulating Th2 cells were positively correlated with total daily dose of inhaled corticosteroid. *ICS*, inhaled corticosteroid.

### Peripheral blood cell production of IL-2 associates with type 2 inflammation

The propensity for high IL-2 production was also related to the degree of type 2 inflammation. Supernatants from patients with high IL-2 following activation of their peripheral blood cells contained more IL-13 (Table 1) and flow cytometry staining of whole blood showed these patients had a higher proportion of CD4^+^ T cells and Th2 cells (CD4^+^ CRTh2^+^ T cells as a proportion of total white blood cells; Table 1). While the association between Th2 cells and IL-2 production was not significant (r = 0.430, *p* = 0.0752), Th2 cells were correlated with total daily dose of inhaled corticosteroid (Fig. 1C, r = 0.583, *p* = 0.011).

### The IL-2-GC axis regulates survival of Th2 cells

Patients with a high proportion of circulating Th2 cells were also taking more inhaled GC (Fig. 1C). GCs induce apoptosis and cell death of various T cell subsets, though to our knowledge no studies have investigated their effect on human Th2 cells. Therefore, we assessed *in vitro* differentiated primary Th2 cell response to DEX (10^−6^ M, 48 hours). Interestingly, we failed to see any increase in Annexin V^+^ cells (Fig. 2A), indicating there was no effect on apoptosis and/or cell death. This was surprising since this level of DEX was able to induce apoptosis in an immortalized T cell line (CCRF CEM, Fig. 2B). A major difference between these two experiments was that the primary Th2 cells were cultured in IL-2 (5 ng/ml). Considering that patients with high IL-2 production had more Th2 cells (Table 1), we next assessed whether titrating the level of this growth factor would alter Th2 cell sensitivity to GC-induced apoptosis. When Th2 cells were cultured with DEX and lower concentrations of IL-2 a significant increase in Annexin V^+^ cells was observed (Fig. 2C). Indeed, Th2 cells showed sensitivity to 10-fold less DEX (10^−7^ M) when cultured with low IL-2 (1.25ng/ml; Fig. 2D) compared to high IL-2 (5ng/ml; Fig. 2C).

**Figure 2.**
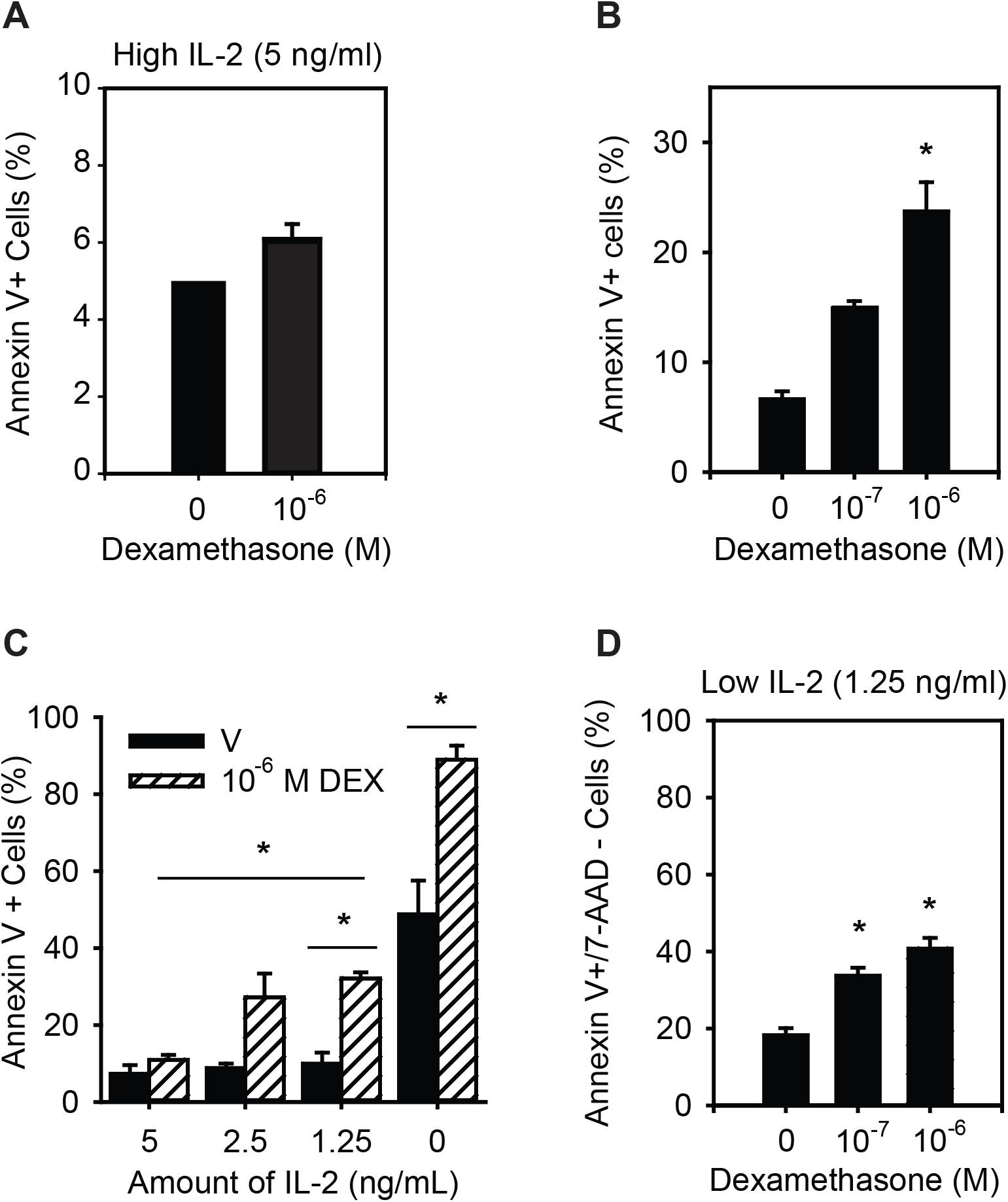
*IL-2 influences Th2 cell sensitivity to GC-induced apoptosis **A)*** Th2 cells cultured in 5ng/ml of IL-2 did not exhibit an increase in Annexin V^+^ cells in response to DEX (10^−6^ M; n = 4). ***B)*** CCRF-CEM cells, an immortalized T cell line, showed an increase in Annexin V^+^ cells in response to DEX (10^−6^ M; n = 4). ***C)*** Titration of IL-2 revealed that the proportion of Th2 cells staining positive for Annexin V increased significantly with lower concentrations of IL-2 (n = 3). ***D)*** Th2 cells cultured in low IL-2 (1.25 ng/ml) exhibited a significant increase in apoptosis Annexin V+/7AAD-cells and responded to 10-fold less DEX (n = 3). Data are presented as mean ± standard error (SE). *V*, vehicle; *DEX*; Dexamethasone; \**p* < 0.05 *vs* vehicle (*i.e*., no DEX).

Apoptosis is regulated by the expression of pro- and anti-apoptotic genes of the BCL-2 family, with the balance between these referred to as the BCL-2 rheostat (38, 39). Since IL-2 induces the anti-apoptotic gene BCL-2 (40) and GC induces the pro-apoptotic gene BIM (41), we assessed expression of these factors. Th2 cells cultured in high IL-2 had more BCL-2 and less BIM mRNA compared to Th2 cells cultured in low IL-2 (Fig. 3A). The ratio of BCL-2 to BIM (BCL-2:BIM) was significantly higher in Th2 cells cultured in high *vs* low IL-2 (Fig. 3B) and remained higher following DEX treatment (Fig. 3C), suggesting its involvement in the observed resistance to GC-induced apoptosis (Fig. 2C). Indeed, when the data for all three DEX concentrations were combined (10^−8^ – 10^−6^ M), the BCL-2:BIM ratio of Th2 cells cultured with high *vs* low IL-2 was significantly higher (Fig. 3D).

**Figure 3.**
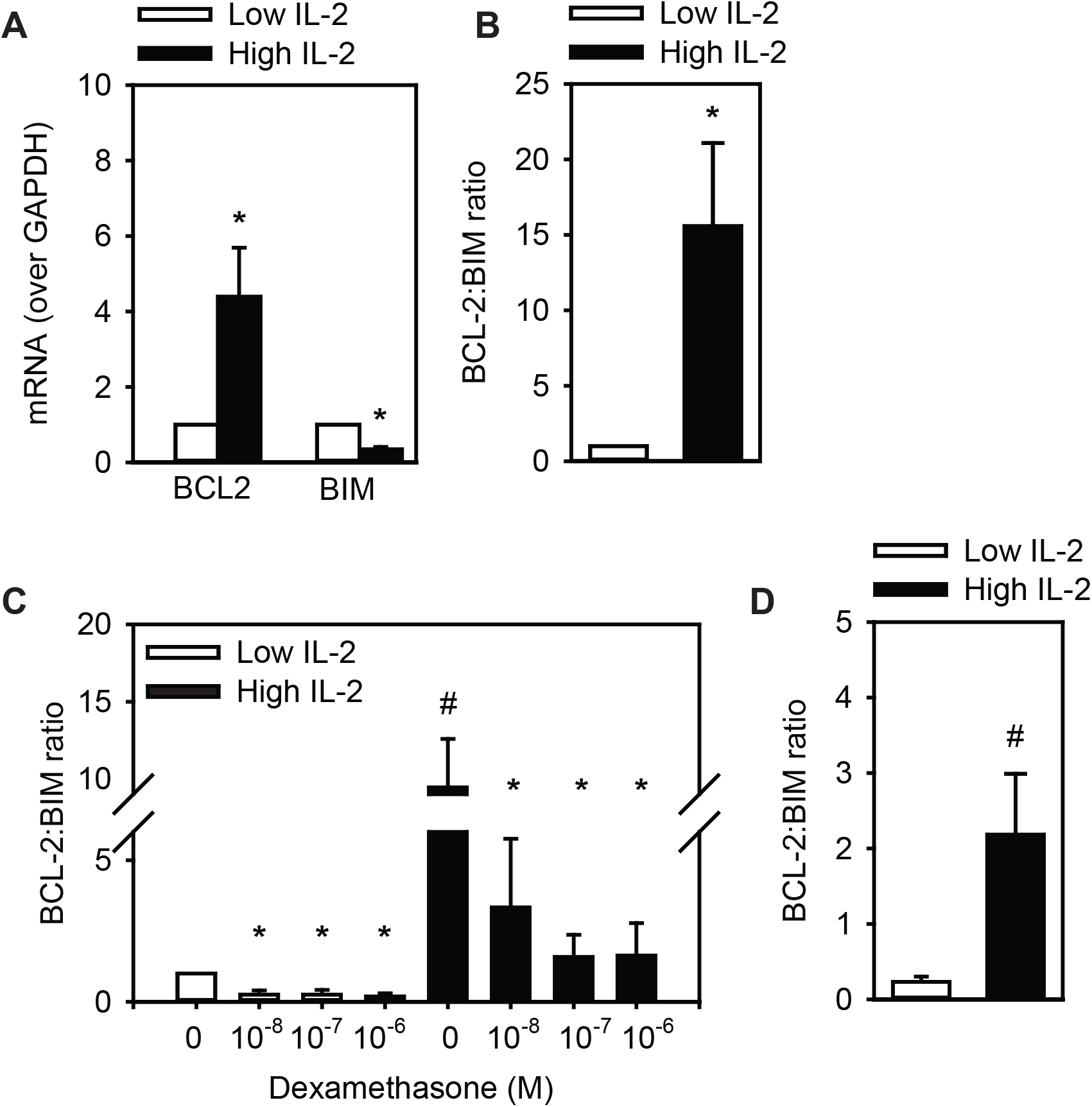
*IL-2 influences the BCL-2 Rheostat **A)*** mRNA for BCL-2 was elevated and BIM was reduced when Th2 cells were cultured in high IL-2 (5 ng/ml) compared to low IL-2 (1.25 ng/ml), ***B)*** leading to an increase in BCL-2:BIM ratio (n = 7). ***C)*** The BCL-2:BIM ratio remains strong (above 1) even after DEX treatment (24 hours, n = 3). ***D)*** Th2 cells cultured in high IL-2 and DEX had greater BCL-2:BIM ratios than low IL-2 and DEX (10^−8^ – 10^−6^ M combined, n = 9). Data are presented as mean ± standard error (SE). *DEX;* Dexamethasone; \**p* < *0.05 vs* no DEX (for each IL-2 concentration); #*p*<*0.05*, high *vs* low IL-2.

### The IL-2-GC axis regulates suppression of type 2 cytokines

Since patients exhibiting high IL-2 production from peripheral blood cells also produced more IL-13 (Table 1), we assessed whether IL-2 directly regulates IL-13 expression. We found that Th2 cells cultured with high IL-2 (24 hours) expressed significantly more IL-13 mRNA compared to those cultured with low IL-2 (Fig. 4A), despite there being no difference in Th2 cell numbers (Fig. 4B).

**Figure 4.**
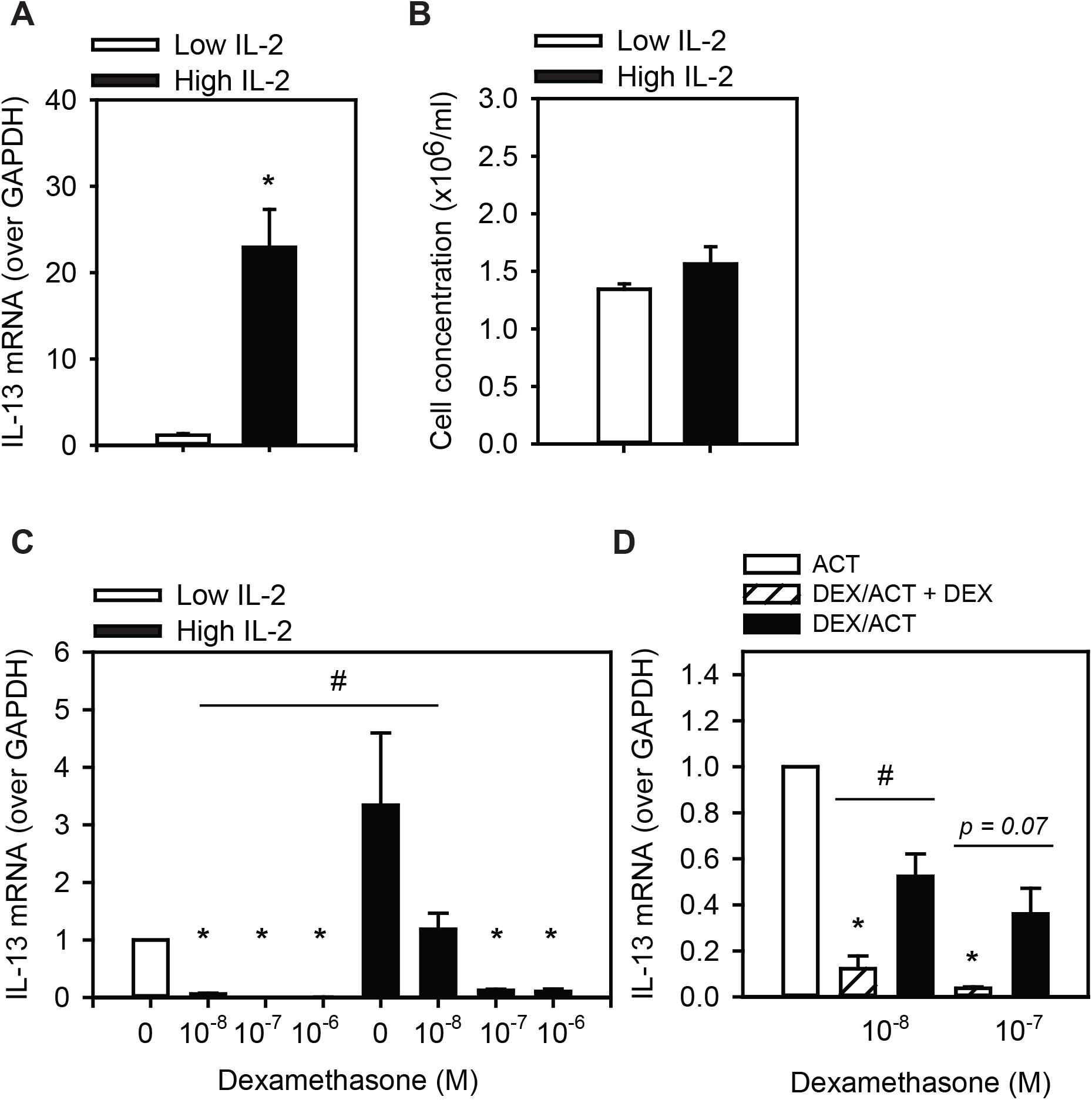
*IL-2 dampens GC-mediated suppression of IL-13 **A)*** Th2 cells cultured with high IL-2 (5 ng/ml) had significantly more IL-13 mRNA compared to those cultured in low IL-2 (1.25 ng/ml; n = 6). ***B)*** Th2 cells cultured in high or low IL-2 showed no difference in cell number (n = 6). ***C)*** The ability of DEX (10^−8^ M) treatment to suppress IL-13 mRNA was significantly less when Th2 cells were cultured in high IL-2 *vs* low IL-2 (n = 3). ***D)*** Continual exposure to DEX was required for suppression of IL-13, since Th2 cells pre-treated with DEX (24 hours) and then activated (ACT) with PMA/ionomycin (24 hours) in the absence of DEX (DEX/ACT) had significantly more IL-13 mRNA than Th2 cells receiving DEX both as a pre-treatment and during activation (DEX/ACT + DEX; n = 3). Data are presented as mean ± standard error (SE). *DEX;* Dexamethasone; \**p* < *0.05 vs* no DEX (for each IL-2 concentration); #*p*<*0.05*, high *vs* low IL-2.

To assess the effect of IL-2 on the ability of GC to suppress type 2 cytokines, we cultured Th2 cells with varying concentrations of IL-2 and DEX. When Th2 cells were cultured in high IL-2 the ability of 10^−8^ M DEX to suppress IL-13 mRNA was significantly less than when cells were cultured with low IL-2 (Fig. 4C), though this difference was not observed at higher DEX concentrations.

To assess the durability of GC suppression following activation, Th2 cells were pretreated with DEX (24hours), washed and activated with mitogen in the presence or absence of DEX (24hours) and high IL-2. Th2 cells receiving DEX as both a pre-treatment and during activation exhibited almost complete suppression of IL-13 mRNA (88 and 96%, respectively), while cells receiving DEX only as a pre-treatment had substantially more IL-13 mRNA following activation (Fig. 4D).

## DISCUSSION

Our study reveals that peripheral blood cell production of IL-2 associates with poor lung function, increased GC usage and heightened type 2 inflammation. *In vitro* experiments showed that maintaining Th2 cells in high IL-2 shifted them toward GC resistance, both at the level of apoptosis and cytokine production. Taken together, these results suggest that an environment of high IL-2 may mediate asthma severity by driving persistence of Th2 cells and their ability to produce type 2 cytokines. Our findings support and extend previous reports indicating a role for IL-2 in GC responsiveness, including Leung *et. al*., which detected more IL-2 mRNA in bronchial alveolar lavage cells of steroid resistant than sensitive asthmatics (42) and Mercado *et. al*. which showed that PBMCs from severe asthmatics produced more IL-2 that non-severe asthmatics (43). Though we did not study this *per se*, severe asthmatics in our population did appear to produce more IL-2. Indeed, the level of IL-2 varied widely and was positively correlated with total daily dose of inhaled steroid, suggesting it may reflect the continuum of GC responsiveness across our population.

IL-2 is a growth factor known to drive proliferation of CD4^+^ T cells (44) and so the higher proportion of Th2 cells in those exhibiting high IL-2 production could be due to having more total CD4^+^ T cells. However, our analyses failed to show a significant difference in the proportion of total CD4^+^ T cells between the high and low IL-2 group, a linear relationship between CD4^+^ T cells and IL-2 (r = 0.354, *p* = 0.150) or differences in growth when Th2 cells were cultured in varying concentrations of IL-2 (1.25 −10 ng/ml). On the other hand, our *in vitro* data did show that Th2 cells cultured in high IL-2 exhibited less GC-induced apoptosis and had more BCL-2 and less BIM mRNA. Furthermore, the BCL-2:BIM ratio remained above 1 (*i.e*., more BCL-2 than BIM) in Th2 cells cultured in high (but not low) IL-2, corresponding with the observed resistance to GC-induced apoptosis. IL-2 induction of BCL-2 is well documented (40, 45). To our knowledge we are the first to report IL-2 supressing BIM expression, though the opposite has been shown – that IL-2 starvation of murine cytotoxic T lymphocytes up-regulated BIM (46). Similarly, Banuelos *et.al*. showed that Th17 cell resistance to GC-induced apoptosis was related to higher expression of BCL-2 (47). Our data highlight the ability of IL-2 to modulate the rheostat, increasing BCL-2 and reducing BIM, to shift Th2 cells toward an anti-apoptotic phenotype. This could suggest that *in vivo* the higher proportion of circulating Th2 cells may be related to the ability of IL-2 to modulate their sensitivity to GC.

Level of IL-2 in the culture media also influenced IL-13 expression, despite no differences in Th2 cell growth, similar to a previous report that IL-2 induced IL-13 expression from antigen-specific Th clones (48). This may be through IL-2 induction of c-Fos (49), a member of the AP-1 complex shown to upregulate IL-13 (50). IL-2 also dampened the capacity of GC to suppress IL-13 transcription. In low IL-2, IL-13 mRNA levels were > 90% suppressed in response to all DEX concentrations, whereas in high IL-2 only a 61% suppression was observed with low DEX. This suggests IL-2 increases GC requirements to achieve cytokine suppression, though simply increasing GC dose may not be effective. Indeed, McKeever *et. al*. reported that asthmatics receiving quadruple their GC dose still exhibited a 45% exacerbation rate in the following year compared to 52% in those that maintained usual GC dose (51). In light of our study, whether this approach could be improved by focusing on ‘low IL-2 producers’, *i.e*. those with a potentially better chance of responding to increased GC, should be assessed.

There are several potential mechanisms underlying the ability to IL-2 to suppress the effects of GC and this heterogeneity contributes to the difficulty of solving the clinical problem of GC resistance (14). For apoptosis, induction of BIM by GC is known to be a direct transcriptional effect as GR binding to regulatory elements within the BIM locus has been shown (52). Therefore, the ability of IL-2 to downregulate BIM could be through IL-2-STAT5 signaling, previously shown to impede GR nuclear translocation (34). In the case of IL-13, less GC-induced inhibition may also be due to IL-2 driving expression of factors that mediate IL-13 transcription such as cFos, a member of the AP-1 complex (49) known to be expressed at higher levels in PBMCs of steroid insensitive asthmatics and to impede GC signaling (53). IL-13 is also regulated by NFκB (54) and IL-2 reverses the GC-mediated induction of IkB, an NFκB inhibitor (54, 55). Yet another mechanism could be through increased activity of the p38MAPK pathway, which can also be activated by IL-2. Indeed, Mercado *et.al*. demonstrated that IL-2/IL-4 treatment of a macrophage cell line (U937) induced GC resistance which could be reversed with an inhibitor of p38MAPK (SB203580) (56). Interestingly, this study also identified two populations of severe asthmatics, based on responsiveness to the p38MAPK inhibitor (56), supporting the view that *in vivo* several mechanisms mediate GC resistance. IL-2 upregulation of GRβ, an inactive form of GR, is another mechanism by which GR signaling may be reduced (57), though the effect was more pronounced in airway than circulating T cells (58) and so some of the complexity surrounding GC resistance may be the result of tissue specific differences.

The strong, super-physiologic activation with PMA and ionomycin resulted in very high levels of IL-2 from patients’ peripheral blood cells, showing their maximal potential for production. In our *in vitro* system we empirically developed a model of high and low IL-2 based on typical concentrations used for Th2 cell cultures (13, 59) as well as functional readouts of resistance/susceptibility to GC-induced apoptosis. Concentrations were lower than observed after *ex vivo* mitogenic activation of patients’ cells, however, are similar to levels found to be produced from *in vivo* antigen-activated T cells (60) and BAL lymphocytes activated using beads coated with αCD2/CD3/CD28 (16) and so may be more physiologically relevant. The source of IL-2 within whole blood could be naïve and/or memory CD4+ T cells (60), Th1 cells (4) or even dendritic cells previously exposure to gram negative bacteria (61). Dendritic cell production would suggest prior infections may influence one’s propensity to produce IL-2 and could even occur *in utero*, as IL-2 promoter methylation at birth was associated with asthma severity in childhood (62). IL-2 genetic variants are also associated with asthma (63, 64) as well as inflammatory bowel disease, type 1 diabetes and multiple sclerosis (63, 65, 66). Together, these data suggest that control of IL-2, whether genetic or epigenetic, may be fundamental to the development of immune disease.

Our finding that suppression of IL-13 required continual exposure to GC, since mRNA levels were double if DEX was received only as a pre-treatment and not also during activation. These data suggest that unless *a)* the dose of GC is sufficient to induce Th2 cell apoptosis and *b)* exposure to GC is continual, suppression of IL-13 is temporary. This could be important clinically, since asthma phenotyping efforts recently proposed a classification based on type 2 cytokine expression (67). However, if a patient is taking a GC dose sufficient to suppress cytokines, but insufficient to eliminate Th2 cells, this could result in misclassification of type 2-low asthma. Using a cell based approach, such as eosinophils and/or Th2 cells, to classify type 2-high asthma may improve identification of the phenotype as well as response to therapies directly target type-2 cells, *i.e*. CRTh2 antagonist (2, 68). Alternative approaches such as BCL-2 inhibitors, which evoke apoptosis and are in use for leukemia (69), also demonstrate promise. In mouse models of both eosinophilic and neutrophilic asthma they were found to be more efficient than steroids to induce granulocyte apoptosis *ex vivo* from patients with severe asthma (70).

In summary, our study shows that IL-2 modulates GC responsiveness of human Th2 cells, supporting both their survival and pro-inflammatory capacity, and suggest that a patient’s potential to produce IL-2 may be a determinant in asthma severity.

## ACKNOWLEDGEMENTS

This study was conducted within the Asthma Center, University of Alberta and so we acknowledge the support of clinical colleagues treating the asthma patients, primarily Drs. Irvin Mayers and Mohit Bhutani. The Authors would also like to thank Angela Hillaby and Miranda Bowen for patient recruitment and sample acquisition; Dr. Cheryl Laratta for developing the patient database; Chad Wu for performing the IL-2 measurements. Funding for this study was from the Canadian Institute of Health Research (LC, HV), Alberta Heritage Foundation for Medical Research (LC), Canadian Lung Association (LC) and the Canadian Allergy Asthma and Immunology Foundation (LS). LC also received funding from a GlaxoSmithKline-Canadian Institute of Health Research Chair in Airway Inflammation (http://ca.gsk.com/en-ca/research/pathfinders-fund/).

## REFERENCES

1. Lötvall J, Akdis CA, Bacharier LB, Bjermer L, Casale TB, Custovic A, Lemanske RF, Wardlaw AJ, Wenzel SE, and Greenberger PA. Asthma endotypes: a new approach to classification of disease entities within the asthma syndrome. J Allergy Clin Immunol. 2011;127(2):355–60.

2. Svenningsen S, and Nair P. Asthma Endotypes and an Overview of Targeted Therapy for Asthma. Front Med (Lausanne). 2017;4(158.

3. Robinson D, Humbert M, Buhl R, Cruz AA, Inoue H, Korom S, Hanania NA, and Nair P. Revisiting Type 2-high and Type 2-low airway inflammation in asthma: current knowledge and therapeutic implications. Clin Exp Allergy. 2017;47(2):161–75.

4. Mosmann TR, and Coffman RL. TH1 and TH2 cells: different patterns of lymphokine secretion lead to different functional properties. Annu Rev Immunol. 1989;7(145–73.

5. Nagata K, Tanaka K, Ogawa K, Kemmotsu K, Imai T, Yoshie O, Abe H, Tada K, Nakamura M, Sugamura K, et al. Selective expression of a novel surface molecule by human Th2 cells in vivo. J Immunol. 1999;162(3):1278–86.

6. Mjosberg JM, Trifari S, Crellin NK, Peters CP, van Drunen CM, Piet B, Fokkens WJ, Cupedo T, and Spits H. Human IL-25- and IL-33-responsive type 2 innate lymphoid cells are defined by expression of CRTH2 and CD161. Nat Immunol. 2011;12(11):1055–62.

7. Nagakumar P, Denney L, Fleming L, Bush A, Lloyd CM, and Saglani S. Type 2 innate lymphoid cells in induced sputum from children with severe asthma. J Allergy Clin Immunol. 2016;137(2):624–6 e6.

8. Smith SG, Chen R, Kjarsgaard M, Huang C, Oliveria JP, O’Byrne PM, Gauvreau GM, Boulet LP, Lemiere C, Martin J, et al. Increased numbers of activated group 2 innate lymphoid cells in the airways of patients with severe asthma and persistent airway eosinophilia. J Allergy Clin Immunol. 2016;137(1):75–86 e8.

9. Camelo A, Rosignoli G, Ohne Y, Stewart RA, Overed-Sayer C, Sleeman MA, and May RD. IL-33, IL-25, and TSLP induce a distinct phenotypic and activation profile in human type 2 innate lymphoid cells. Blood Adv. 2017;1(10):577–89.

10. Islam SA, and Luster AD. T cell homing to epithelial barriers in allergic disease. Nat Med. 2012;18(5):705–15.

11. Mojtabavi N, Dekan G, Stingl G, and Epstein MM. Long-lived Th2 memory in experimental allergic asthma. J Immunol. 2002;169(9):4788–96.

12. Woodland DL, and Kohlmeier JE. Migration, maintenance and recall of memory T cells in peripheral tissues. Nat Rev Immunol. 2009;9(3):153–61.

13. Shamji MH, Temblay JN, Cheng W, Byrne SM, Macfarlane E, Switzer AR, Francisco NDC, Olexandra F, Jacubczik F, Durham SR, et al. Antiapoptotic serine protease inhibitors contribute to survival of allergenic TH2 cells. J Allergy Clin Immunol. 2017.

14. Barnes PJ. Glucocorticosteroids: current and future directions. Br J Pharmacol. 2011;163(1):29–43.

15. Christodoulopoulos P, Cameron L, Durham S, and Hamid Q. Molecular pathology of allergic disease. II: Upper airway disease. J Allergy Clin Immunol. 2000;105(2 Pt 1):211–23.

16. Kaur M, Reynolds S, Smyth LJ, Simpson K, Hall S, and Singh D. The effects of corticosteroids on cytokine production from asthma lung lymphocytes. Int Immunopharmacol. 2014;23(2):581–4.

17. Jee YK, Gilmour J, Kelly A, Bowen H, Richards D, Soh C, Smith P, Hawrylowicz C, Cousins D, Lee T, et al. Repression of interleukin-5 transcription by the glucocorticoid receptor targets GATA3 signaling and involves histone deacetylase recruitment. J Biol Chem. 2005;280(24):23243–50.

18. Chung KF, Wenzel SE, Brozek JL, Bush A, Castro M, Sterk PJ, Adcock IM, Bateman ED, Bel EH, Bleecker ER, et al. International ERS/ATS guidelines on definition, evaluation and treatment of severe asthma. Eur Respir J. 2014;43(2):343–73.

19. Nair P, Pizzichini MM, Kjarsgaard M, Inman MD, Efthimiadis A, Pizzichini E, Hargreave FE, and O’Byrne PM. Mepolizumab for prednisone-dependent asthma with sputum eosinophilia. N Engl J Med. 2009;360(10):985–93.

20. Haldar P, Brightling CE, Hargadon B, Gupta S, Monteiro W, Sousa A, Marshall RP, Bradding P, Green RH, Wardlaw AJ, et al. Mepolizumab and exacerbations of refractory eosinophilic asthma. NEngl J Med. 2009;360(10):973–84.

21. Corren J, Lemanske RF, Hanania NA, Korenblat PE, Parsey MV, Arron JR, Harris JM, Scheerens H, Wu LC, Su Z, et al. Lebrikizumab treatment in adults with asthma. NEngl J Med. 2011;365(12):1088–98.

22. Ortega HG, Liu MC, Pavord ID, Brusselle GG, FitzGerald JM, Chetta A, Humbert M, Katz LE, Keene ON, Yancey SW, et al. Mepolizumab Treatment in Patients with Severe Eosinophilic Asthma. New England Journal of Medicine. 2014;371(13):1198–207.

23. Distelhorst CW. Recent insights into the mechanism of glucocorticosteroid-induced apoptosis. Cell Death Differ. 2002;9(1):6–19.

24. Zhang X, Moilanen E, and Kankaanranta H. Enhancement of human eosinophil apoptosis by fluticasone propionate, budesonide, and beclomethasone. Eur J Pharmacol. 2000;406(3):325–32.

25. Woolley KL, Gibson PG, Carty K, Wilson AJ, Twaddell SH, and Woolley MJ. Eosinophil apoptosis and the resolution of airway inflammation in asthma. Am J Respir Crit Care Med. 1996;154(1):237–43.

26. Marchetti MC, Di Marco B, Cifone G, Migliorati G, and Riccardi C. Dexamethasone-induced apoptosis of thymocytes: role of glucocorticoid receptor-associated Src kinase and caspase-8 activation. Blood. 2003;101(2):585–93.

27. Brinkmann V, and Kristofic C. Regulation by corticosteroids of Th1 and Th2 cytokine production in human CD4+ effector T cells generated from CD45RO- and CD45RO+ subsets. J Immunol. 1995;155(7):3322–8.

28. Tischner D, Theiss J, Karabinskaya A, van den Brandt J, Reichardt SD, Karow U, Herold MJ, Lühder F, Utermöhlen O, and Reichardt HM. Acid sphingomyelinase is required for protection of effector memory T cells against glucocorticoid-induced cell death. J Immunol. 2011;187(9):4509–16.

29. Banuelos J, and Lu NZ. A gradient of glucocorticoid sensitivity among helper T cell cytokines. Cytokine Growth Factor Rev. 2016;31(27–35.

30. Palikhe NS, Laratta C, Nahirney D, Vethanayagam D, Bhutani M, Vliagoftis H, and Cameron L. Elevated levels of circulating CD4(+) CRTh2(+) T cells characterize severe asthma. Clin Exp Allergy. 2016;46(6):825–36.

31. Fajt ML, Gelhaus SL, Freeman B, Uvalle CE, Trudeau JB, Holguin F, and Wenzel SE. Prostaglandin D(2) pathway upregulation: relation to asthma severity, control, and TH2 inflammation. J Allergy Clin Immunol. 2013;131(6):1504–12.

32. Liao W, Lin JX, and Leonard WJ. IL-2 family cytokines: new insights into the complex roles of IL-2 as a broad regulator of T helper cell differentiation. Curr Opin Immunol. 2011;23(5):598–604.

33. Ross SH, and Cantrell DA. Signaling and Function of Interleukin-2 in T Lymphocytes. Annu Rev Immunol. 2018;36(411–33.

34. Goleva E, Kisich KO, and Leung DY. A role for STAT5 in the pathogenesis of IL-2-induced glucocorticoid resistance. J Immunol. 2002;169(10):5934–40.

35. Nieto MA, and López-Rivas A. IL-2 protects T lymphocytes from glucocorticoid-induced DNA fragmentation and cell death. J Immunol. 1989;143(12):4166–70.

36. Moffatt MF, Gut IG, Demenais F, Strachan DP, Bouzigon E, Heath S, von Mutius E, Farrall M, Lathrop M, Cookson WOCM, et al. A large-scale, consortium-based genomewide association study of asthma. NEngl J Med. 2010;363(13):1211–21.

37. MacLean Scott E, Solomon LA, Davidson C, Storie J, Palikhe NS, and Cameron L. Activation of Th2 cells downregulates CRTh2 through an NFAT1 mediated mechanism. PLoS One. 2018;13(7):e0199156.

38. Ploner C, Rainer J, Niederegger H, Eduardoff M, Villunger A, Geley S, and Kofler R. The BCL2 rheostat in glucocorticoid-induced apoptosis of acute lymphoblastic leukemia. Leukemia. 2008;22(2):370–7.

39. Volkmann N, Marassi FM, Newmeyer DD, and Hanein D. The rheostat in the membrane: BCL-2 family proteins and apoptosis. Cell Death Differ. 2014;21(2):206–15.

40. Ahmed NN, Grimes HL, Bellacosa A, Chan TO, and Tsichlis PN. Transduction of interleukin-2 antiapoptotic and proliferative signals via Akt protein kinase. Proc Natl Acad Sci U S A. 1997;94(8):3627–32.

41. Heidari N, Miller AV, Hicks MA, Marking CB, and Harada H. Glucocorticoid-mediated BIM induction and apoptosis are regulated by Runx2 and c-Jun in leukemia cells. Cell Death Dis. 2012;3(e349.

42. Leung DY, Martin RJ, Szefler SJ, Sher ER, Ying S, Kay AB, and Hamid Q. Dysregulation of interleukin 4, interleukin 5, and interferon gamma gene expression in steroid-resistant asthma. J Exp Med. 1995;181(1):33–40.

43. Mercado N, Hakim A, Kobayashi Y, Meah S, Usmani OS, Chung KF, Barnes PJ, and Ito K. Restoration of corticosteroid sensitivity by p38 mitogen activated protein kinase inhibition in peripheral blood mononuclear cells from severe asthma. PLoS One. 2012;7(7):e41582.

44. Morgan DA, Ruscetti FW, and Gallo R. Selective in vitro growth of T lymphocytes from normal human bone marrows. Science. 1976;193(4257):1007–8.

45. Akbar AN, Borthwick NJ, Wickremasinghe RG, Panayoitidis P, Pilling D, Bofill M, Krajewski S, Reed JC, and Salmon M. Interleukin-2 receptor common gamma-chain signaling cytokines regulate activated T cell apoptosis in response to growth factor withdrawal: selective induction of anti-apoptotic (bcl-2, bcl-xL) but not proapoptotic (bax, bcl-xS) gene expression. Eur J Immunol. 1996;26(2):294–9.

46. Stahl M, Dijkers PF, Kops GJ, Lens SM, Coffer PJ, Burgering BM, and Medema RH. The forkhead transcription factor FoxO regulates transcription of p27Kip1 and Bim in response to IL-2. J Immunol. 2002;168(10):5024–31.

47. Banuelos J, Shin S, Cao Y, Bochner BS, Morales-Nebreda L, Budinger GR, Zhou L, Li S, Xin J, Lingen MW, et al. BCL-2 protects human and mouse Th17 cells from glucocorticoid-induced apoptosis. Allergy. 2016;71(5):640–50.

48. Hashimoto T, Kobayashi N, Kajiyama Y, Kaminuma O, Suko M, and Mori A. IL-2-induced IL-13 production by allergen-specific human helper T cell clones. Int Arch Allergy Immunol. 2006;140 Suppl 1(51–4.

49. Kawahara A, Minami Y, Miyazaki T, Ihle JN, and Taniguchi T. Critical role of the interleukin 2 (IL-2) receptor gamma-chain-associated Jak3 in the IL-2-induced c-fos and c-myc, but not bcl-2, gene induction. Proc Natl Acad Sci U S A. 1995;92(19):8724–8.

50. Lorentz A, Klopp I, Gebhardt T, Manns MP, and Bischoff SC. Role of activator protein 1, nuclear factor-kappaB, and nuclear factor of activated T cells in IgE receptor-mediated cytokine expression in mature human mast cells. J Allergy Clin Immunol. 2003;111(5):1062–8.

51. McKeever T, Mortimer K, Wilson A, Walker S, Brightling C, Skeggs A, Pavord I, Price D, Duley L, Thomas M, et al. Quadrupling Inhaled Glucocorticoid Dose to Abort Asthma Exacerbations. N Engl J Med. 2018;378(10):902–10.

52. Jing D, Bhadri VA, Beck D, Thoms JA, Yakob NA, Wong JW, Knezevic K, Pimanda JE, and Lock RB. Opposing regulation of BIM and BCL2 controls glucocorticoid-induced apoptosis of pediatric acute lymphoblastic leukemia cells. Blood. 2015;125(2):273–83.

53. Lane SJ, Adcock IM, Richards D, Hawrylowicz C, Barnes PJ, and Lee TH. Corticosteroid-resistant bronchial asthma is associated with increased c-fos expression in monocytes and T lymphocytes. J Clin Invest. 1998;102(12):2156–64.

54. Pahl A, Zhang M, Kuss H, Szelenyi I, and Brune K. Regulation of IL-13 synthesis in human lymphocytes: implications for asthma therapy. Br J Pharmacol. 2002;135(8):1915–26.

55. Xie H, Seward RJ, and Huber BT. Cytokine rescue from glucocorticoid induced apoptosis in T cells is mediated through inhibition of IkappaBalpha. Mol Immunol. 1997;34(14):987–94.

56. Mercado N, To Y, Kobayashi Y, Adcock IM, Barnes PJ, and Ito K. p38 mitogen-activated protein kinase-gamma inhibition by long-acting beta2 adrenergic agonists reversed steroid insensitivity in severe asthma. Mol Pharmacol. 2011;80(6):1128–35.

57. Leung DY, Hamid Q, Vottero A, Szefler SJ, Surs W, Minshall E, Chrousos GP, and Klemm DJ. Association of glucocorticoid insensitivity with increased expression of glucocorticoid receptor beta. J Exp Med. 1997;186(9):1567–74.

58. Hamid QA, Wenzel SE, Hauk PJ, Tsicopoulos A, Wallaert B, Lafitte JJ, Chrousos GP, Szefler SJ, and Leung DY. Increased glucocorticoid receptor beta in airway cells of glucocorticoid-insensitive asthma. Am JRespir Crit Care Med. 1999;159(5 Pt 1):1600–4.

59. Ray JP, Staron MM, Shyer JA, Ho PC, Marshall HD, Gray SM, Laidlaw BJ, Araki K, Ahmed R, Kaech SM, et al. The Interleukin-2-mTORc1 Kinase Axis Defines the Signaling, Differentiation, and Metabolism of T Helper 1 and Follicular B Helper T Cells. Immunity. 2015;43(4):690–702.

60. Sojka DK, Bruniquel D, Schwartz RH, and Singh NJ. IL-2 secretion by CD4+ T cells in vivo is rapid, transient, and influenced by TCR-specific competition. J Immunol. 2004;172(10):6136–43.

61. Granucci F, Vizzardelli C, Pavelka N, Feau S, Persico M, Virzi E, Rescigno M, Moro G, and Ricciardi-Castagnoli P. Inducible IL-2 production by dendritic cells revealed by global gene expression analysis. Nat Immunol. 2001;2(9):882–8.

62. Curtin JA, Simpson A, Belgrave D, Semic-Jusufagic A, Custovic A, and Martinez FD. Methylation of IL-2 promoter at birth alters the risk of asthma exacerbations during childhood. Clin Exp Allergy. 2013;43(3):304–11.

63. Matesanz F, Fedetz M, Leyva L, Delgado C, Fernandez O, and Alcina A. Effects of the multiple sclerosis associated −330 promoter polymorphism in IL2 allelic expression. JNeuroimmunol. 2004;148(1-2):212–7.

64. Christensen U, Haagerup A, Binderup HG, Vestbo J, Kruse TA, and Borglum AD. Family based association analysis of the IL2 and IL15 genes in allergic disorders. Eur J Hum Genet. 2006;14(2):227–35.

65. Festen EA, Goyette P, Scott R, Annese V, Zhernakova A, Lian J, Lefèbvre C, Brant SR, Cho JH, Silverberg MS, et al. Genetic variants in the region harbouring IL2/IL21 associated with ulcerative colitis. Gut. 2009;58(6):799–804.

66. Fichna M, Zurawek M, Fichna P, Ziółkowska-Suchanek I, Januszkiewicz D, and Nowak J. Polymorphic variant at the IL2 region is associated with type 1 diabetes and may affect serum levels of interleukin-2. Mol Biol Rep. 2013;40(12):6957–63.

67. Woodruff PG, Modrek B, Choy DF, Jia G, Abbas AR, Ellwanger A, Koth LL, Arron JR, and Fahy JV. T-helper type 2-driven inflammation defines major subphenotypes of asthma. Am JRespir Crit Care Med. 2009;180(5):388–95.

68. White C, Wright A, and Brightling C. Fevipiprant in the treatment of asthma. Expert Opin Investig Drugs. 2018;27(2):199–207.

69. Yogarajah M, and Stone RM. A concise review of BCL-2 inhibition in acute myeloid leukemia. Expert Rev Hematol. 2018;11(2):145–54.

70. Tian BP, Xia LX, Bao ZQ, Zhang H, Xu ZW, Mao YY, Cao C, Che LQ, Liu JK, Li W, et al. Bcl-2 inhibitors reduce steroid-insensitive airway inflammation. J Allergy Clin Immunol. 2017;140(2):418–30.

